# Self-amplifying RNA SARS-CoV-2 lipid nanoparticle vaccine induces equivalent preclinical antibody titers and viral neutralization to recovered COVID-19 patients

**DOI:** 10.1101/2020.04.22.055608

**Authors:** Paul F. McKay, Kai Hu, Anna K. Blakney, Karnyart Samnuan, Clément R. Bouton, Paul Rogers, Krunal Polra, Paulo J.C. Lin, Christopher Barbosa, Ying Tam, Robin J. Shattock

## Abstract

The spread of the SARS-CoV-2 into a global pandemic within a few months of onset motivates the development of a rapidly scalable vaccine. Here, we present a self-amplifying RNA encoding the SARS-CoV-2 spike protein encapsulated within a lipid nanoparticle as a vaccine and demonstrate induction of robust neutralization of a pseudo-virus, proportional to quantity of specific IgG and of higher quantities than recovered COVID-19 patients. These data provide insight into the vaccine design and evaluation of immunogenicity to enable rapid translation to the clinic.

## MAIN

The unprecedented and rapid spread of SARS-CoV-2 into a global pandemic, with the current estimated number of confirmed cases >2.2 million people,^1^ has motivated the need for a rapidly producible and scalable vaccine. Coronaviruses are positive-sense, single stranded RNA viruses that cause disease pathology ranging from the common cold to pneumonia.^2,3^ Despite being listed on the WHO blueprint priority list, there are currently no licensed vaccines for SARS or MERS.^4^ However, previous studies have elucidated the need to stabilize coronavirus spike proteins in their pre-fusion conformation in order to serve as a vaccine immunogen.^5^

Self-amplifying RNA (saRNA) encapsulated in lipid nanoparticles (LNP) is a highly relevant platform for producing vaccines in the context of a global pandemic as it’s possible to encode any antigen of interest^6,7^ and requires a minimal dose compared to messenger RNA (mRNA).^8^ The first RNA therapeutic, which is formulated in LNP, was approved in 2018 and has set the precedent for clinical safety of LNP-formulated RNA.^9^

Here, we compare the immunogenicity of saRNA encoding a pre-fusion stabilized SARS-CoV-2 spike protein encapsulated in LNP in a preclinical murine model to the immune response generated by a natural infection in recovered COVID-19 patients. We characterize both the humoral and cellular response as well as the neutralization capacity of a pseudotyped SARS-CoV-2 virus.

## RESULTS

After confirming expression of the pre-fusion stabilized SARS-CoV-2 spike protein *in vitro* (Supplementary Figure 1), mice were immunized with saRNA encoding the SARS-CoV-2 spike protein encapsulated in LNP with doses ranging from 0.01 to 10 μg (Figure 1a). Mice received two injections, one month apart, and electroporated plasmid DNA (pDNA) was used as a positive control while saRNA encoding the rabies glycoprotein (RABV) in pABOL was used as a negative control. After 6 weeks, we observed remarkably high quantities of SARS-CoV-2 specific IgG in mouse sera in a dose-responsive manner, ranging from 10^5^-10^6^ ng/mL (Figure 1b). The groups that received doses of 10 and 1 μg of saRNA LNP were significantly higher than the mice that received 10 μg of electroporated pDNA, with p=0.0036 and 0.0020, respectively. All of the saRNA LNP-vaccinated mice, even the 0.01 μg group, had higher quantities of SARS-CoV-2 specific IgG compared to patients that had recovered from COVID-19, which had a mean titer of 10^3^ ng/mL and a range of 10^1^-10^5^ ng/mL. Importantly both the pDNA and saRNA LNP immunizations induced a Th1-biased response in mice (Supplementary Figure 2).

**Figure 1.**
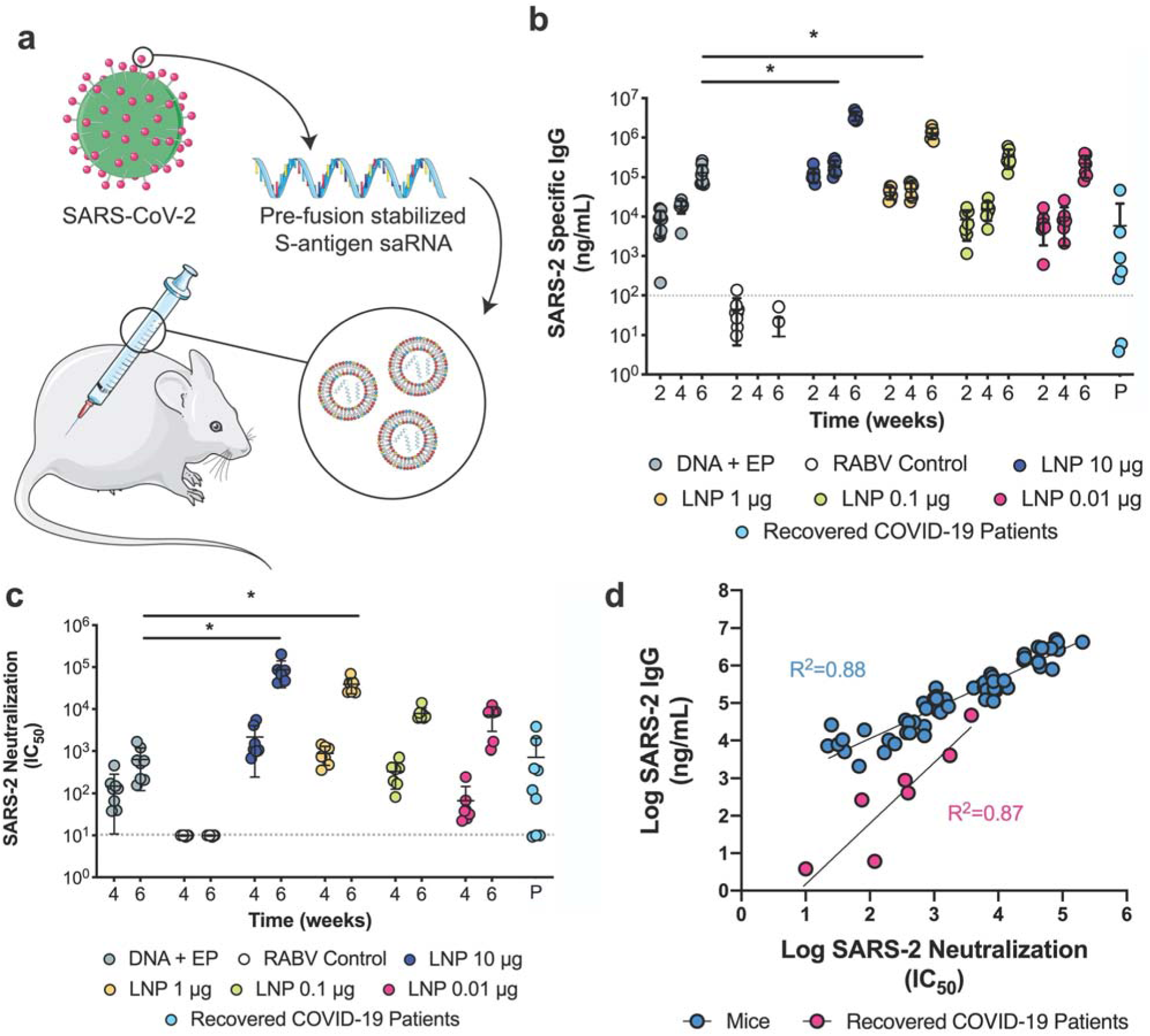
Antibody quantification and neutralization of a SARS-CoV-2 saRNA vaccinated mice compared to COVID-19 recovered patients. a) Schematic of vaccination of BALB/c mice with saRNA encoding pre-fusion stabilized spike protein in LNP, b) SARS-CoV-2 specific IgG responses in mice vaccinated with doses of LNP-formulated saRNA ranging from 0.01-10 μg of saRNA with n=7 and COVID-19 recovered patients with n=9, c) SARS-CoV-2 pseudotyped virus neutralization of sera from BALB/c mice vaccinated with doses of LNP-formulated saRNA ranging from 0.01-10 μg of saRNA with n=7 and COVID-19 recovered patients with n=9, d) Correlation between SARS-CoV-2-specific IgG and SARS-CoV-2 neutralization IC_50_ for vaccinated mice (n=7) and recovered COVID-19 patients (n=9). Electroporated pDNA (DNA + EP) was used as a positive control while saRNA encoding the rabies glycoprotein (RABV) in pABOL was used as a negative control (RABV control). * indicates significance of p<0.05.

We then sought to characterize how antibodies generated by immunization compared to those generated by a natural SARS-CoV-2 infection as far as capacity to neutralize a SARS-CoV-2 pseudotyped virus (Figure 1c). We observed highly efficient viral neutralization that varied in a linear dose-dependent manner for the mice vaccinated with saRNA LNP, with IC_50_ values ranging from 5×10^3^ to 10^5^. The groups that received 10 or 1 μg of saRNA LNP were significantly higher than the electroporated pDNA positive control group, both with p<0.0001. Comparison to the IC_50_ values of recovered COVID-19 patients, which had an average IC_50_ of 10^3^, revealed that even the lowest dose of saRNA LNP (0.01 μg) in mice induced higher SARS-CoV-2 neutralization than a natural infection in humans.

We then determined if there is a correlation between the quantity of SARS-CoV-2 specific IgG and SARS-CoV-2 neutralization IC_50_ for both vaccinated mice and patients who have recovered from COVID-19. Both mice and patients have positive correlations between antibody level and viral neutralization, with R^2^= 0.88 and 0.87 and p<0.0001 and =0.0007, respectively, indicating that high antibody titers enable more efficient viral neutralization. We also tested the sera of vaccinated mice and recovered patients against other pseudotyped viruses, including SARS-CoV, MERS-CoV and 229E-CoV (Supplementary Figure 3), and observed slight neutralization of SARS-CoV by vaccinated mice sera, but otherwise no cross-reactivity.

We also characterized the cellular response and induction of systemic cytokines in response to vaccination with saRNA LNP (Figure 2). We observed that splenocytes from vaccinated mice re-stimulated with a library of SARS-CoV-2 peptides yielded remarkably high IFN-Y secretion as quantified by ELISpot (Figure 2a). The saRNA LNP groups that received 0.01-10 μg ranged from 1,000-2,600 SFU/10^6^ splenocytes, and the 1 and 10 μg groups were significantly higher than the EP pDNA positive control group, with p=0.0016 and 0.0078, respectively. The re-stimulated splenocyte secretions were also characterized with a panel of cytokines (Supplementary Figure 4), with notable increases in GM-CSF, IL-10, IL-12, IL-17a, IL-21, IL-4, IL-5, IL-6, TNF-α, IP-10, MIP-1β and RANTES.

**Figure 2.**
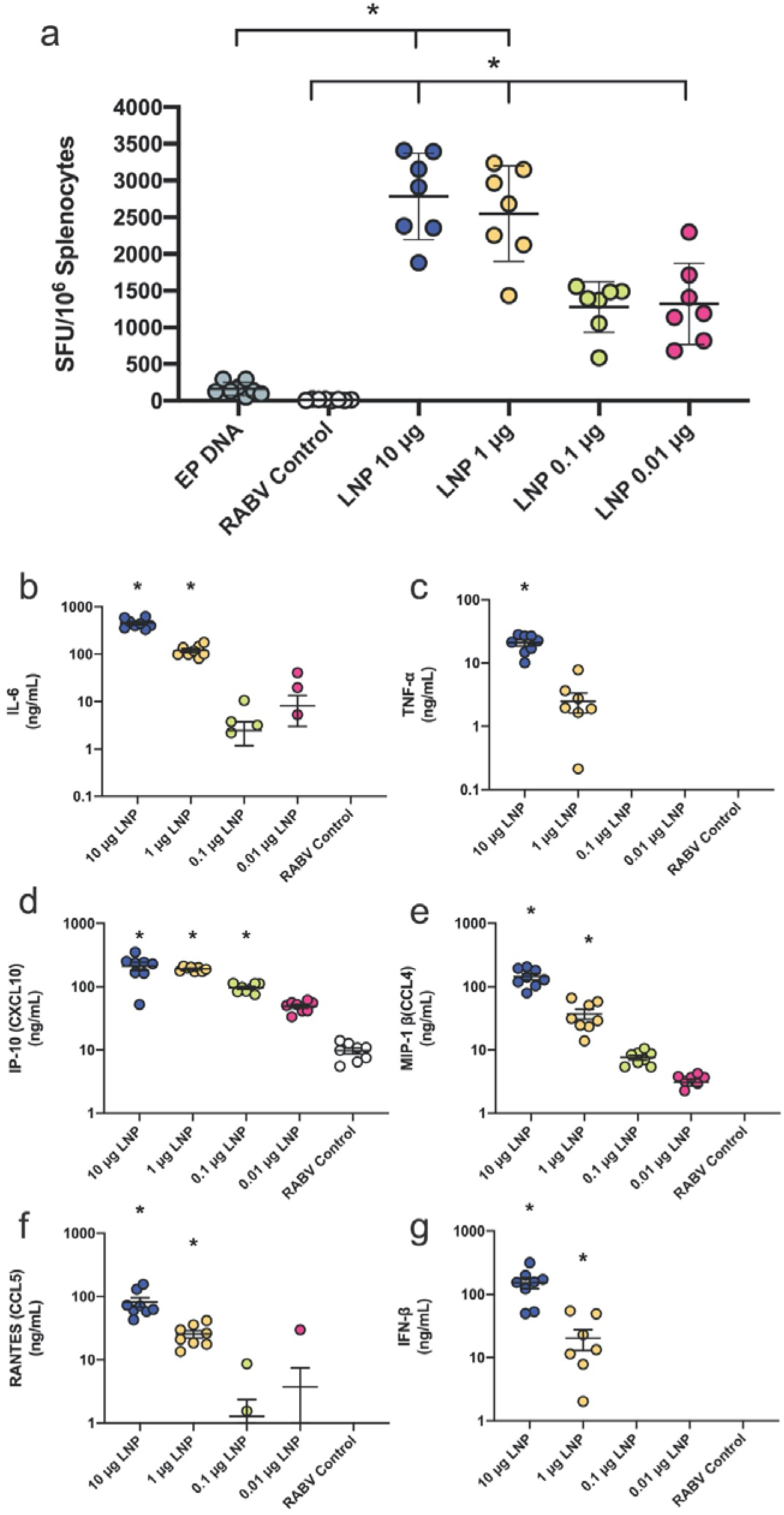
Cellular and secreted cytokine responses to a SARS-CoV-2 saRNA LNP vaccine. a) Quantification of IFN-Y splenocytes upon restimulation with SARS-CoV-2 peptides, expressed as spot forming units (SFU) per 10^6^ cells with n=7. Electroporated pDNA (EP pDNA) was used as a positive control while saRNA encoding the rabies glycoprotein (RABV) in pABOL was used as a negative control (RABV control). b-g) Cytokine profile in sera of mice 4 hours after vaccination with SARS-CoV-2 LNP vaccine with n=7. Remaining cytokines can be found in Supplementary Figure 5.

We further characterized the immune response by assessing the systemic cytokine response 4 hours after injection with LNP-formulated saRNA LNP (Figure 2b-g). The groups that received 10 and 1 μg of saRNA LNP had enhanced levels of IL-6, MIP-1β, RANTEs, IFN-β and IP-10 in the sera compared to the RABV control group, indicating that the LNP formulation enables the immunogenicity of the saRNA. Data from the complete cytokine panel is presented in Supplementary Figure 5.

## DISCUSSION

Here we characterized the immunogenicity of a SARS-CoV-2 saRNA LNP vaccine compared to the immune response of a natural infection in COVID-19 recovered patients. We observed that two saRNA LNP immunizations induced remarkably high SARS-CoV-2 specific IgG antibodies in mice, with quantities that were superior to both EP pDNA and natural infection in humans, that were able to efficiently neutralize a pseudotyped virus. We also observed that the saRNA LNP vaccine induces a robust cellular response, which is partially enabled by the potent LNP formulation.

We observed that the saRNA-encoded pre-fusion stabilized spike protein of SARS-CoV-2 used in these studies is highly immunogenic, yielding antibody titers >10^6^ ng/mL (Figure 1), which is superior to what others have reported for subunit vaccines for the SARS, MERS and SARS-2 coronaviruses.^10^ Furthermore, we observed higher antibody titers, viral neutralization (IC_50_) and cellular response for LNP-formulated saRNA than electroporated pDNA, which we postulate is due to the potent LNP used in these studies, as previous comparisons between polyplex-formulated saRNA and EP pDNA have yielded similar immunogenicity.^8^ This is highly useful for translation as it means a potent LNP-formulated saRNA vaccine can be injected with a widely accepted syringe and needle, and does not require electroporation instrumentation, which we envision will enable more widespread vaccination to curb the spread of SARS-CoV-2.

The saRNA LNP vaccine presented in these studies elicited robust antibody and cellular responses, with a Th1 bias that we hypothesize will enable immunogenicity in humans. Ongoing studies are being carried out to characterize the potential for antibody dependent enhancement (ADE) of SARS-CoV-2 as has been observed for SARS and MERS,^11,12^ but the role of this phenomena in vaccine-induced immunity is not yet fully understood. Overall, we believe that these data inform the antigen design, formulation and preclinical evaluation of immunogenicity that will enable rapid translation of a SARS-CoV-2 vaccine to the clinic trials.

## METHODS

### Vectors

We used a plasmid vector to synthesize a self-amplifying RNA (saRNA) replicon, based on a Trinidad donkey Venezuelan equine encephalitis virus strain (VEEV) alphavirus genome. The viral structural proteins driven from the sub-genomic promoter were replaced by the surface ‘spike’ glycoprotein of the novel severe acute respiratory syndrome coronavirus 2 (SARS-CoV-2): GenBank accession number: QHD43416.1 with some modifications.^13^ The pre-fusion state of the spike glycoprotein was stabilized by proline substitutions of K968 and V969. We synthesized oligonucleotide fragments encoding the SARS-CoV-2 gene using GeneArt strings (Thermo Fisher Scientific) and assembled into the plasmid vector with NEB HiFi assembly (New England BioLabs). An expression plasmid expressing the same pre-fusion stabilized full length transmembrane protein used the pcDNA3.1 backbone and was directly synthesized and cloned into the vector by GeneArt (Thermo Fisher Scientific). A plasmid that expressed a soluble pre-fusion version was directly synthesized and cloned into the pcDNA3.1 backbone vector by GeneArt (Thermo Fisher Scientific). This soluble version ends at glutamine Q1208 of the pre-fusion modified QHD43416.1 gene sequence followed by a GGGGSGGGGS linker, a T4 fibritin (foldon) trimerization motif, a further GGGGSGGGGS linker, the Myc tag, a GSGSGS linker and finally an 8xHIS tag to enable purification of the soluble recombinant protein. The RABV control saRNA was based on the Pasteur strain: GenBank accession number: NP_056796.1 with the F318V amino acid substitution to reduce glycoprotein binding to the neurotrophin receptor (p75NTR), a natural ligand.

### Recombinant soluble SARS-CoV-2 S expression and purification

The plasmid expressing the soluble pre-fusion version of SARS-CoV-2 S was used to produce the recombinant protein using the FreeStyle™ 293 Expression System (Thermo Fisher Scientific), according to the manufacturer’s instructions. Conditioned medium was clarified by centrifugation and protein was sequentially purified by a HisTrap HP column and a HiPrep 16/60 Sephacryl S-300 HR size exclusion chromatography (SEC) column (both from GE Healthcare). Purified protein was first analyzed by Native-PAGE and Western blot, and then filtered through a 0.22 μm membrane, aliquoted and stored at −80 °C.

### *In Vitro* Transcription of RNA

Self-amplifying RNA encoding the pre-fusion stabilized SARS-CoV-2 was produced using *in vitro* transcription. pDNA was transformed into *E. coli* (New England BioLabs, UK), cultured in 100 mL of Luria Broth (LB) with 100 μg/mL carbenicillin (Sigma Aldrich, UK). Plasmid was purified using a Plasmid Plus MaxiPrep kit (QIAGEN, UK) and the concentration and purity was measured on a NanoDrop One (ThermoFisher, UK). pDNA was linearized using MluI for 3h at 37°C. Uncapped *in vitro* RNA transcripts were produced using 1 μg of linearized DNA template in a MEGAScript™ reaction (Ambion, UK) for 2h at 37°C, according to the manufacturer’s protocol. Transcripts were then purified by overnight LiCl precipitation at −20°C, centrifuged at 14,000 RPM for 20 min at 4°C to pellet, washed with 70% EtOH, centrifuged at 14,000 RPM for 5 min at 4°C and resuspended in UltraPure H_2_O (Ambion, UK). Purified transcripts were capped using the ScriptCap™ Cap 1 Capping System Kit (CellScript, WI, USA) for 2h at 37°C, according to the manufacturer’s protocol. Capped transcripts were purified by LiCl precipitation as described above, resuspended in RNA storage buffer (10 mM HEPES, 0.1 mM EDTA, and 100 mg/mL trehalose) and stored at −80°C until further use.

### Cell Culture & saRNA Transfection

HEK293T/17 cells (ATCC) were cultured in complete Dulbecco’s Modified Eagle’s Medium (DMEM) (Gibco, Thermo Fisher Scientific) containing 10 % fetal bovine serum (FBS, Gibco, Thermo Fisher Scientific), 1 % L-glutamine and 1 % penicillin-streptomycin (Thermo Fisher Scientific) at 37°C, 5% CO₂. Cells were plated in a 12-well plate at a density of 0.75 × 10^6^ cells per well 48 h prior to transfection. Lipofectamine MessengerMAX (Thermo Fisher Scientific) was used according to the manufacturer’s instructions for the transfection of SARS-CoV-2 saRNA.

### Flow Cytometry

Twenty-four hours post transfection, cells were harvested and resuspended in 1 mL of FACS buffer (PBS + 2.5 % FBS) at a concentration of 1 × 10^7^ cells /mL. One hundred microliters of the resuspended cells was added to a FACS tube and stained with 50 µL of Live/Dead Fixable Aqua Dead Cell Stain (Thermo Fisher Scientific) at a 1:400 dilution on ice for 20 min. Cells were then washed with 2.5 mL of FACS buffer and centrifuged at 1750 RPM for 7 min. After centrifugation, cells were stained with 2.5 µg of a SARS-CoV spike protein polyclonal antibody (PA1-41165, Thermo Fisher Scientific) for 30 min on ice before washing with 2.5 mL of FACS buffer and centrifuging at 1750 RPM for 7 min. Cells were then stained with 0.4 µg of FITC goat anti-rabbit IgG (BD Pharmigen) for 30 min on ice. After incubation, cells were washed with 2.5 mL of FACS buffer, centrifuged at 1750 RPM for 7 min and resuspended with 250 µL of PBS. Cells were fixed with 250 µL of 3 % paraformaldehyde for a final concentration of 1.5 %. Samples were analyzed on a LSRForterssa (BD Biosciences) with FACSDiva software (BD Biosciences). Data were analyzed using FlowJo Version 10 (FlowJo LLC).

### Formulation of saRNA

saRNA was encapsulated in LNP using a self-assembly process in which an aqueous solution of saRNA at pH=4.0 is rapidly mixed with an ethanolic lipid mixture.^14^ LNP used in this study were similar in composition to those described previously^15,16^, which contain an ionizable cationic lipid (proprietary to Acuitas)/phosphatidylcholine/cholesterol/PEG-lipid. The proprietary lipid and LNP composition are described in US patent US10,221,127. They had a mean hydrodynamic diameter of ∼75nm with a polydispersity index of <0.1 as measured by dynamic light scattering using a Zetasizer Nano ZS (Malvern Instruments Ltd, Malvern, UK) instrument and an encapsulation efficiency of >90% LNP.

RABV control group was formulated with 8 kDa pABOL at a ratio of polymer to RNA of 45:1 (w/w) using the titration method as previously described.^7^

### Animals and immunizations

BALB/c mice aged 6-8 weeks old were placed into groups of n = 7 or 8. Animals were handled and procedures were performed in accordance with the terms of a project license granted under the UK Home Office Animals (Scientific Procedures) Act 1986. All the procedures and protocols used in this study were approved by an animal ethical committee, the Animal Welfare and Ethical Review Body (AWERB). Groups of mice were injected intramuscularly (IM; quadriceps) with a 50 µL of vaccine saRNA formulations. For animals that were vaccinated with pDNA, 10 µg of pDNA was injected in 50 µL PBS followed by electroporation (EP) using 5-mm electrodes using an ECM 830 square-wave electroporation system (BTX) (pulses: 100 V of positive and negative polarity at 1 pulse/s, 50 ms pulse). Animals were immunized at week 0, boosted with a second vaccination at week 4 and euthanized using a Schedule 1 method at week 6, at which time the spleens were removed and processed to single cells for use in assays. Serum samples were collected at two-week intervals.

### Recovered COVID-19 patient samples

Serum samples were donated to the Communicable Diseases Research Tissue Bank, Section of Virology, Imperial College London, following written informed consent, by patients who had been infected with SARS-CoV-2. The tissue bank is approved by the National Research Ethics Service, South Central Committee Oxford C (Ref 15/SC/0089).

### IFN-γ ELISpots

Assessment of the IFN-γ T cell response was performed using the Mouse IFN-γ ELISpot^PLUS^ kit (Mabtech) following the manufacturer’s instructions. Briefly, anti-IFN-γ pre-coated plates were blocked with DMEM + 10% FBS for at least 30min, then cells were added at 2.5×10^5^ cells/well for negative control (media only) and SARS-CoV-2 peptide pools (15-mers overlapping by 11; JPT Peptides) (1 µg/mL) in 200 µL final volume per well. The positive control wells contained 5×10^4^ cells/well in 200 µL final volume per well with 5 µg/mL of ConA. Plates were incubated overnight at 5% CO2, 37ºC incubator and developed as per the manufacturer’s protocol. Once dried, plates were read using the AID ELISpot reader ELR03 and AID ELISpot READER software (Autoimmun Diagnostika GmbH).

### Antigen-specific Ig ELISA

The antigen-specific IgG, IgG1 and IgG2a titres in mouse sera were assessed by a semi-quantative ELISA as previously described.^17^ In brief, MaxiSorp high binding ELISA plates (Nunc) were coated with 100 μL/well of 1 μg/mL recombinant SARS-CoV-2 protein in PBS. For the standard IgG/IgG1/IgG2a, 3 columns on each plate were coated with 1:1000 dilution each of goat anti-mouse Kappa and Lambda light chains (Southern Biotech). After overnight incubation at 4ºC, the plates were washed 4 times with PBS-Tween 20 0.05% (v/v) and blocked for 1 h at 37ºC with 200 μL/well blocking buffer (1% BSA (w/v) in PBS-Tween 20 0.05%(v/v)). The plates were then washed and the diluted samples or a 5-fold dilution series of the standard IgG (or IgG1 or IgG2) added using 50 μL/well volume. Plates were incubated for 1 h at 37ºC, then washed and secondary antibody added at 1:2000 dilution in blocking buffer (100 μL/well) using either anti-mouse IgG-HRP, anti-mouse IgG1-HRP or anti-mouse IgG2a-HRP (Southern Biotech). After incubation and washes, plates were developed using 50 μL/well SureBlue TMB (3,3’, 5,5’-tetramethylbenzidine) substrate and the reaction stopped after 5 min with 50 μL/well stop solution (Insight Biotechnologies). The absorbance was read on a Versamax Spectrophotometer at 450 nm (BioTek Industries).

### Neutralization assay

A HIV-pseudotyped luciferase-reporter based system was used to assess the neutralization ability of sera from vaccinated animals and recovered patients against SARS-CoV, SARS-CoV-2, MERS-CoV and 229E-CoV, as previously described with modifications.^18,19^ In brief, CoV S-pseudotyped viruses were produced by co-transfection of 293T/17 cells with a HIV-1 *gag-pol* plasmid (pCMV-Δ8.91, a kind gift from Prof. Julian Ma, St George’s University of London), a firefly luciferase reporter plasmid (pCSFLW, a kind gift from Prof. Julian Ma, St George’s University of London) and a plasmid encoding the S protein of interest (pSARS-CoV-S, pSARS-CoV2-S, pMERS-CoV-S or p229E-CoV-S) at a ratio of 1:1.5:1. Virus-containing medium was clarified by centrifugation and filtered through a 0.45 μm membrane 72h after transfection, and subsequently aliquoted and stored at −80 °C. For the neutralization assay, heat-inactivated sera were first serially diluted and incubated with virus for 1 h, and then the serum-virus mixture was transferred into wells pre-seeded Caco2 cells. After 48h, cells were lysed and luciferase activity was measured using Bright-Glo Luciferase Assay System (Promega). The IC_50_ neutralization was then calculated using GraphPad Prism (version 8.4).

### Cytokine measurement in splenocytes and sera

Splenoctyes isolated from each individual mouse were plated into round bottom 96 well plates (1 × 10^6^ per well in a 200 uL total volume) and cultured for 7 days with media alone, 5 ug / mL SARS-CoV-2 recombinant protein or 5 ug / mL ConA as a positive control. For the sera samples, mice were bled 4h after injection with SARS-CoV-2 LNP vaccine or control RABV vaccine and sera were collected. The cytokine response in each well was quantified with a custom 25-plex ProcartaPlex Immunoassay (ThermoFisher Scientific, UK) on a Bio-Plex 200 System (Bio-Rad), according to the manufacturer’s instructions.

### Statistical analysis

Graphs and statistics were prepared in GraphPad Prism (version 8.4). Statistical differences were analyzed using either a two-way ANOVA adjusted for multiple comparisons or a Kruskal-Wallis test adjusted for multiple comparisons, with p<0.05 used to indicate significance.

## Data availability

Raw data is available upon reasonable request from Imperial College London.

## ACKNOWLEDGEMENTS

We gratefully acknowledge Graham Cooke, Rachael Quinlan, Charlotte Short, and Carolina Rosadas de Oliveira for providing the samples from recovered COVID-19 patients, and Jonathan Yeow for providing the polymers used in these studies. AKB is supported by a Marie Skłodowska Curie Individual Fellowship funded by the European Commission H2020 (No. 794059). PFM, KH, KS, CRB, KP, PR and RJS are funded by the Department of Health and Social Care using UK Aid funding and is managed by the Engineering and Physical Sciences Research Council (EPSRC, grant number: EP/R013764/1, note: the views expressed in this publication are those of the author(s) and not necessarily those of the Department of Health and Social Care). This work was supported in part by the NIHR Biomedical Research Centre of Imperial College Healthcare NHS Trust. We also acknowledge Dormeur Investment Services Ltd for providing funds to purchase equipment used in these studies.

## AUTHOR CONTRIBUTIONS

PFM and RJS conceptualized the antigen design and designed the studies. PFM, KS and KH designed and performed *in vitro* experiments. PFM, AKB, KH and KS performed *in vivo* studies, aided by CRB, KP, and PR. PL, CB and YT designed and prepared the saRNA LNP. AKB analysed the data and wrote the manuscript with help from PFM, KH and KS, and constructive feedback and editing from CRB, KP, PR and RJS.

