## Supplementary Information for "Self-amplifying RNA SARS-CoV-2 lipid nanoparticle vaccine induces equivalent preclinical antibody titers and viral neutralization to recovered COVID-19 patients"

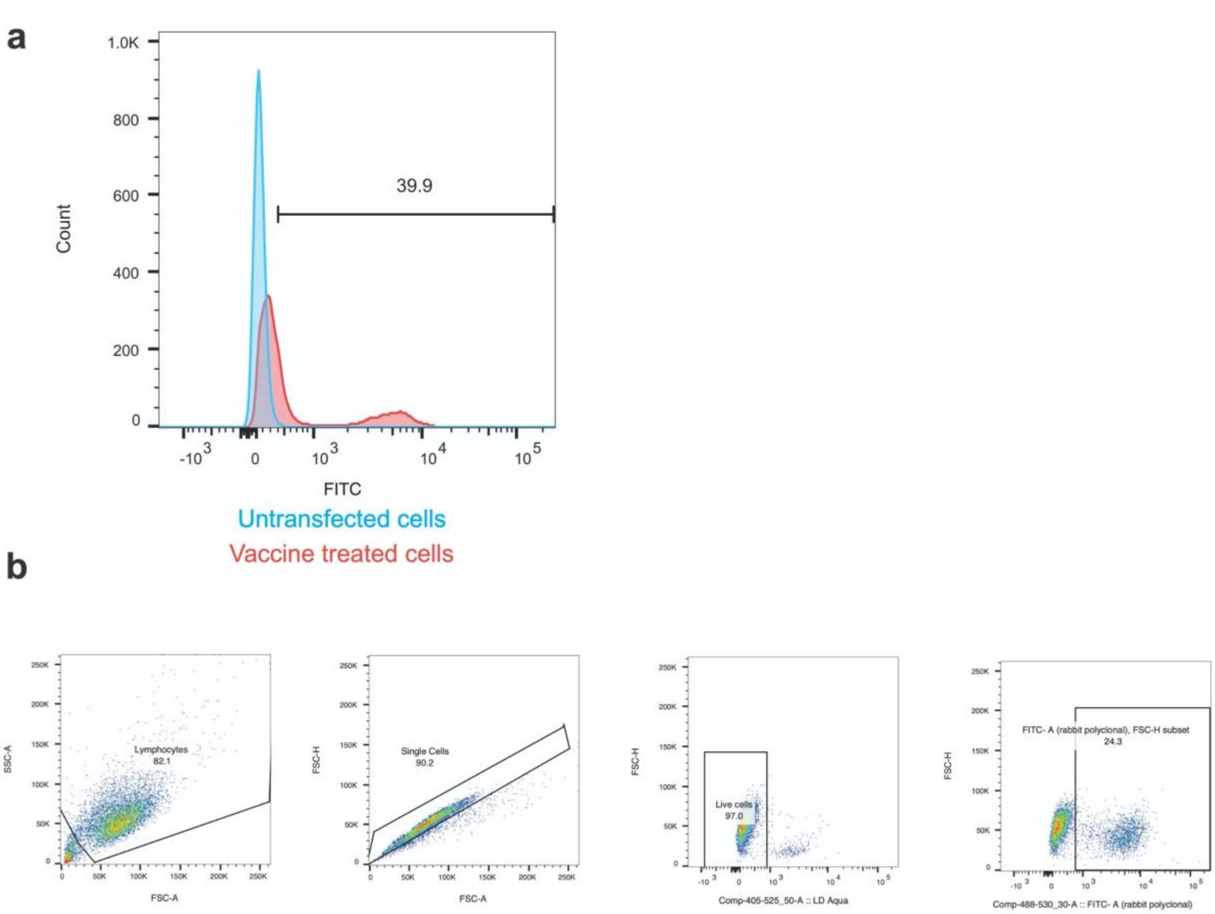


**Supplementary Figure 1**. Flow cytometry of HEK293T.17 cells transfected with vaccine expressing membrane bound SARS-CoV-2 spike protein.

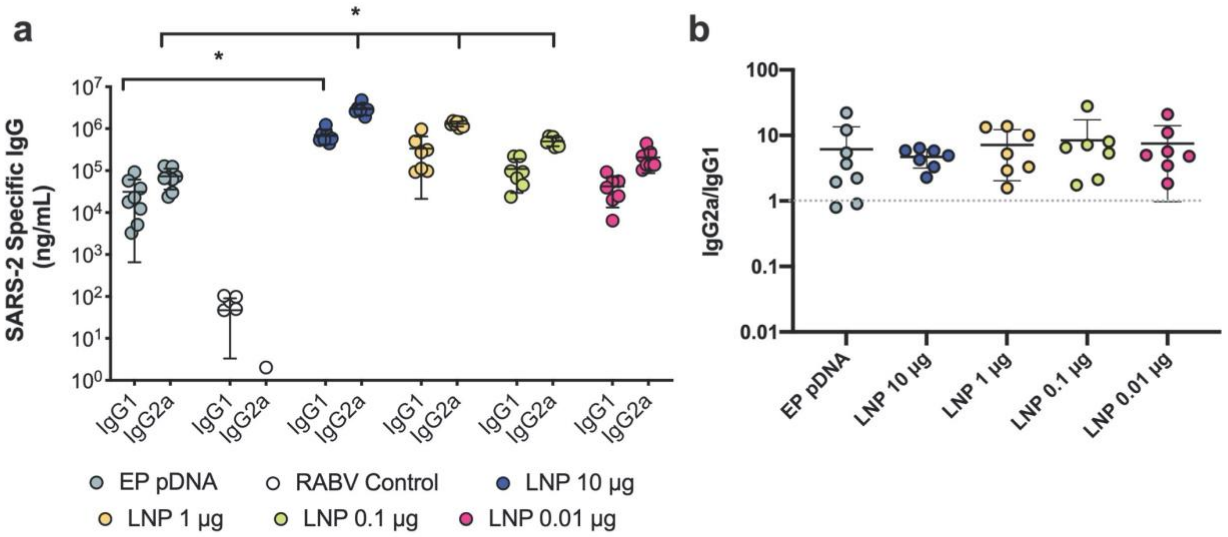


**Supplementary Figure 2**. Th1/Th2 skew in response to SARS-CoV-2 saRNA LNP vaccine. a) IgG1 and IgG2a responses in mice vaccinated with doses of LNP-formulated saRNA ranging from 0.01-10 μg of saRNA with n=7, b) Th1/Th2 skewing responses in mice vaccinated with doses of LNP-formulated saRNA ranging from 0.01-10 μg of saRNA with n=7, and 10 μg of electroporated pDNA (EP pDNA) with n=8.

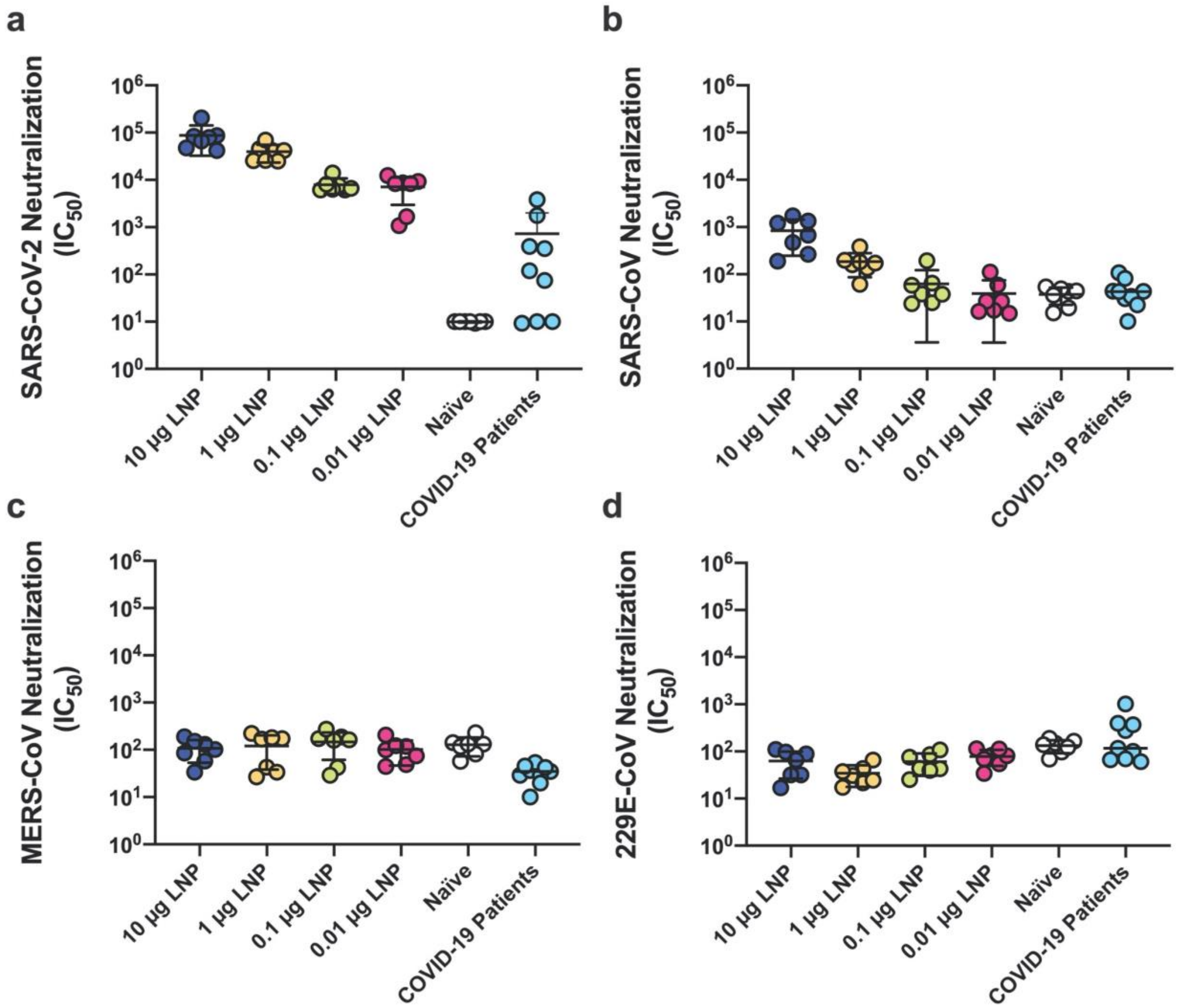


**Supplementary Figure 3**. Pseudotyped virus neutralization of sera from BALB/c mice vaccinated with LNP formulations and human patients after recovery from COVID-19.

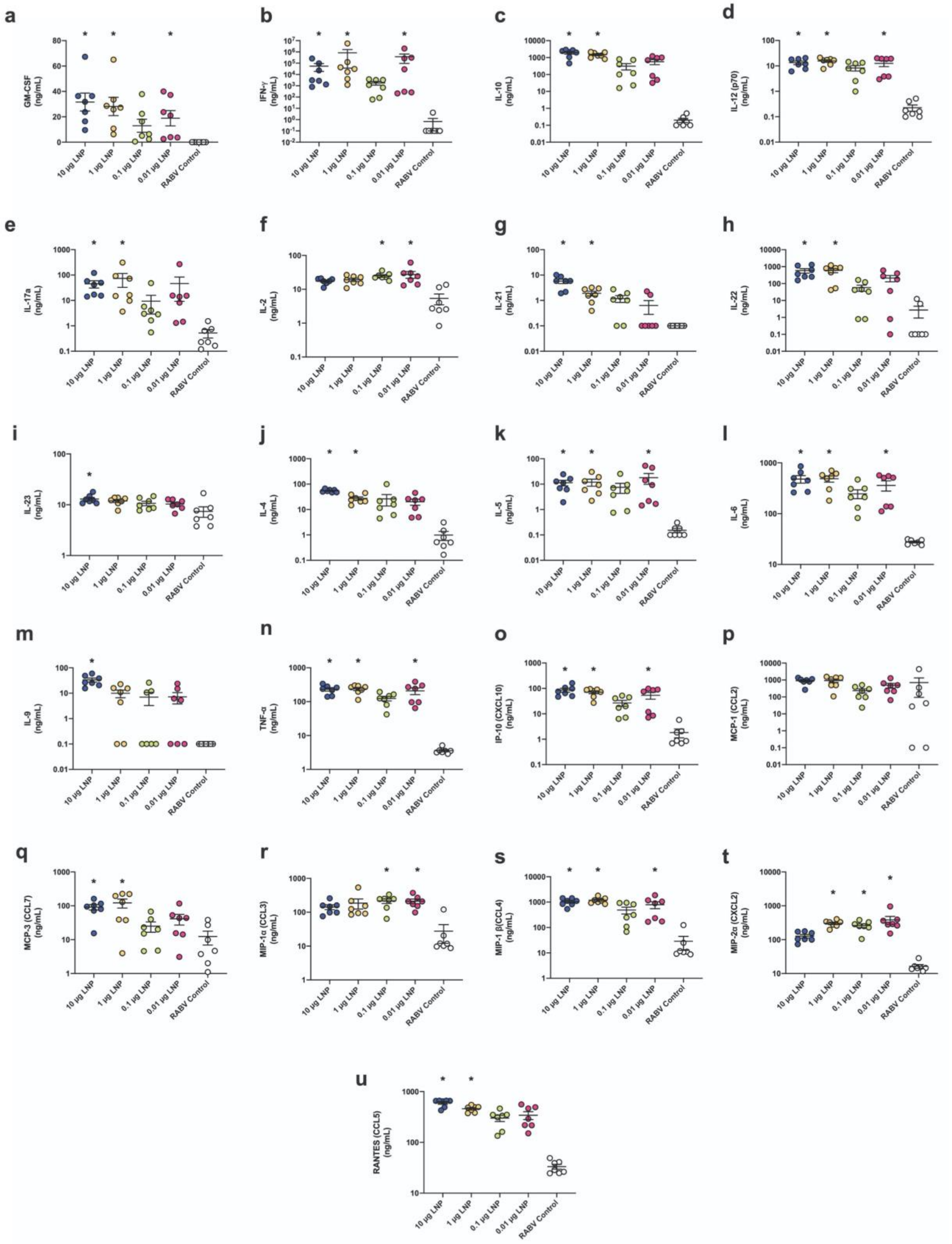


**Supplementary Figure 4**. Cytokine response of re-stimulated splenocytes of vaccinated mice with recombinant SARS-CoV-2 protein.

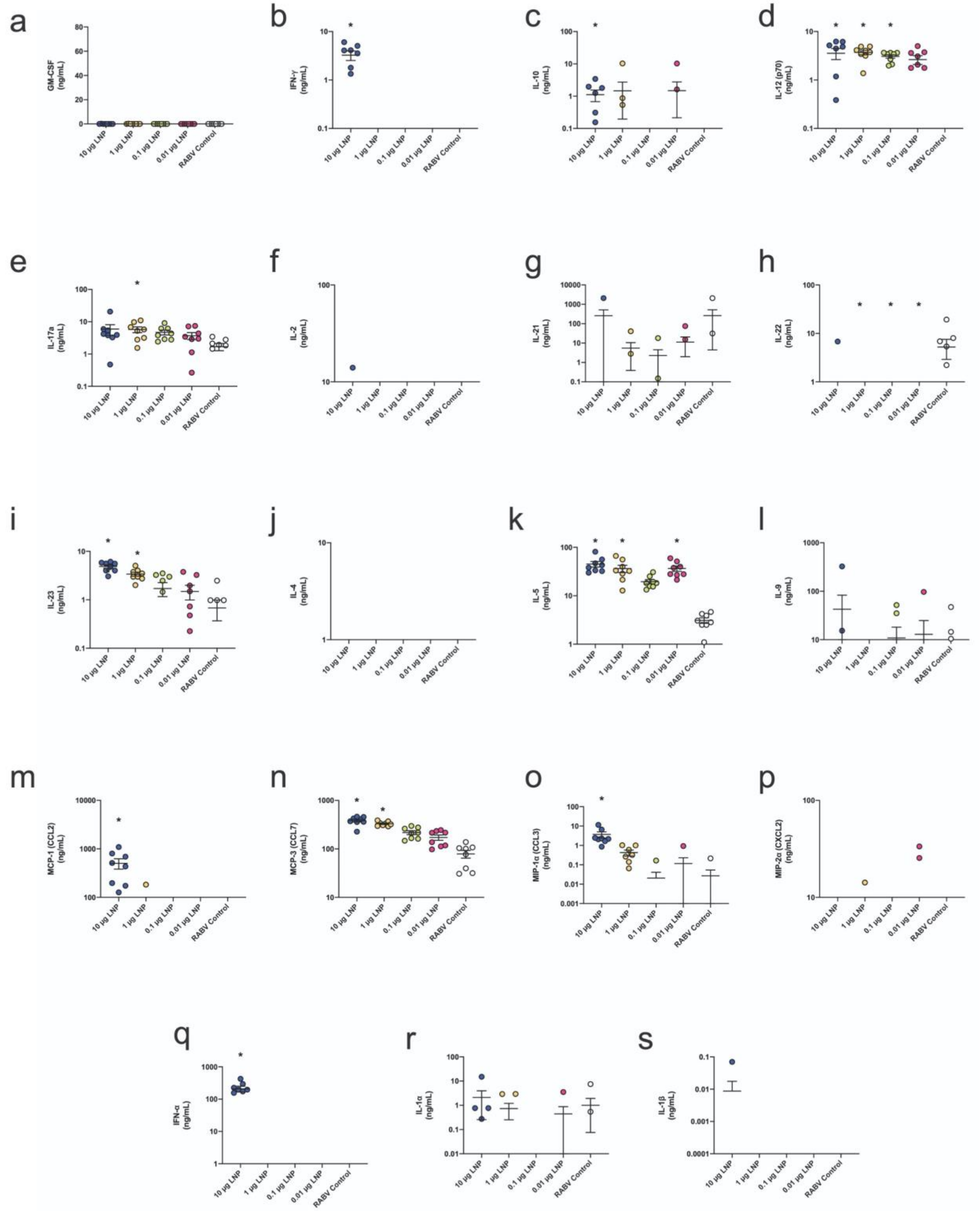


**Supplementary Figure 5**. Cytokine response in the sera of mice 4 hours after vaccination with SARS-CoV-2 LNP vaccine.
